# Innovation in solitary bees is driven by exploration, shyness and activity levels

**DOI:** 10.1101/2019.12.23.884619

**Authors:** Miguel Á. Collado, Randolf Menzel, Daniel Sol, Ignasi Bartomeus

## Abstract

Behavioural innovation is widely considered an important mechanism by which animals respond to novel environmental challenges, including those induced by human activities. Despite its functional and ecological relevance, much of our current understanding of the innovation process comes from studies in vertebrates. Understanding innovation processes in insects has lagged behind partly because they are not perceived to have the cognitive machinery required to innovate. This perception is however challenged by recent evidence demonstrating sophisticated cognitive capabilities in insects despite their small brains. Here, we study the innovation capacity of a solitary bee (*Osmia cornuta*) in the laboratory by exposing naïve individuals to an obstacle removal task. We also studied the underlying cognitive and non-cognitive mechanisms through a battery of experimental tests designed to measure learning, exploration, shyness and activity levels. We found that solitary bees can innovate, with 11 of 29 individuals (38%) being able to solve a new task consisting in lifting a lid to reach a reward. The propensity to innovate was uncorrelated with learning capacities, but increased with exploration, boldness and activity. These results provide solid evidence that non-social insects can innovate, and highlight the importance of interpreting innovation in the light of non-cognitive processes.

## INTRODUCTION

Animals exhibit an extraordinary wide repertoire of behaviours. Bees, for example, have developed a broad repertoire of sophisticated behaviours that facilitate foraging, nesting, navigation, and communication (Roulston & Goodell, 2011) Although the ecological and evolutionary importance of behaviour is widely recognised, our current understanding of how new behaviours emerge is insufficiently understood. Some simple behaviours have a clear genetic basis, and hence may have been acquired through mutation and natural selection. Studies in *Drosophila* show, for example, that a mutation in a single neuropeptide caused several abnormalities on their behavioural circadian rhythms (i.e. biological clocks, Renn et *al*., 1999). However, the accumulation of mutations seems insufficient to understand the emergence of complex behaviours. Rather, the emergence of novel behaviours from more simple cognitive processes require the processing of new knowledge by means of experience to guide decision-making (Dukas, 2008). The emergence of new learnt behaviours is a process known as behavioural innovation (Ramsey et *al.*, 2007, Lefebvre et *al.*, 2004, Reader et *al.*, 2003, Sol 2003).

The concept of innovation has attracted considerable interest of researchers for its broad implications for ecology and evolution (Ramsey et *al*., 2007; Lefebvre et *al*., 2004; Reader, 2003; Sol, 2003). Innovating designates the possibility of constructing plastic behavioural responses to novel ecological challenges, thereby potentially enhancing the fitness of the individual animals when exposed to unusual or novel situations. For instance, evidence is accumulating that innovation abilities enhances the success of animals when introduced to novel environments (Sol et *al*., 2005). By changing the relationship of individuals with the environment, innovative behaviours also have a great potential to influence the evolutionary responses of the population to selective pressures (Lefebvre et *al*., 2004; Reader et *al*., 2016). Hence, in a context of global change, innovative behaviours are considered central to understand how animals will respond to rapid changes induced by human activities.

While innovation is considered one of the main processes behind the emergence of novel behaviours in vertebrates (Reader, 2003; Ramsey et *al*., 2007), the relevance of innovation is currently insufficiently understood in insects. The traditional notion holds that insect behaviour tends to be relatively inflexible and stereotypical, a perception that partially arises from their small brains and less number of neurons than more studied taxa like mammals or birds (Dukas, 2008). Such a belief is however changing as evidence accumulates of unsuspected sophisticated capabilities that transcend basic forms of cognition, including rule learning (Gil et *al*., 2007), numerosity (Chittka et *al*., 1995, Dacke & Srinivasan, 2008), development of novel routes and shortcuts while navigating (Menzel et *al*., 2005) or exploratory learning (Menzel & Giurfa, 2001; Degen et *al*., 2016). The fact that insects exhibits sophisticated cognition suggests that new behaviours may also be commonly acquired through the process of innovation.

Here, we address the critical questions of whether insects are capable of innovate and how they achieve it. We used a solitary common bee —*Osmia cornuta* (Megachilidae)— as a model system to address these questions. While our current understanding of cognition in solitary bees is limited in comparison to that of eusocial species (e.g. Chittka & Thompson, 2009), they are also easy to rear and manipulate in captivity (Jin et *al.* 2014). An advantage of solitary bees is that they can be tested individually for innovative propensity without having to consider the pitfall of separating individuals from the social group. Importantly, solitary bees compose most of the bee fauna and are suffering worldwide population declines associated with rapid human-induced environmental changes (Goulson et *al.*, 2015), posing at risk the essential pollination services that they provide for cultivated crops and wild plants (Ollerton, J, Tarrant, S & Winfree, R 2011). Thus, there is an urgent need to assess whether and how they are capable of innovate to cope with new environmental challenges.

The capacity to innovate is difficult to measure directly (Lefebvre et *al.*, 2004), but one widely adopted approach is the use of problem-solving experiments motivated by a food reward (Bouchard et *al.*, 2007, Griffin et *al.*, 2014). In our experiments, we exposed naïve *O. cornuta* bees to a novel task consisting in lifting a lid to reach a food reward, an assay that mimics the encounter of a new complex flower. Whether or not individuals solve the task and the latency in doing so can be used as measures of innovation performance (Sol et *al*., 2011). Because some bees were capable to innovate, we investigated the underlying mechanisms. We first explored whether the propensity to innovate reflects a domain-general ability to learn. Hence, we related our measures of innovation performance to measures of performance in an associative learning test. Next, we tested the effect of a number of emotional and state-dependent intrinsic features that are suspected to either facilitate or inhibit innovation (Reader et *al.*, 2003, Houston & McNamara, 1999; Sol et *al.*, 2012), including exploration, shyness and activity levels. We finally considered whether problem-solving ability might be explained by sex, an additional intrinsic parameter (Houston & McNamara 1999). In *O. cornuta*, females are more involved in parental activities (e.g. are in charge of all nest provisioning activities) and are typically larger than males (Bosch, 1994). These fundamental differences in the biology and ecology between sexes are expected to affect how they deal with novel challenges, potentially affecting their problem-solving ability.

## MATERIAL AND METHODS

### Study subjects

*Osmia cornuta* cocoons were bought from the company WAB-Mauerbienenzucht (Konstanz, Deutschland) and kept cold at 4C°. Before and during the experiments, cocoons were put in 15 ml falcon tubes in a pitch black environment and kept in an incubator at 26°C for 24-48 hours until the emergence of offspring. In total, 101 females and 42 males were born, and used in the experiments. In order to force bees to walk instead of fly, we anesthetized them with a cold shock treatment and cut their right wings (Crook, 2013).

### Experimental device

We conducted the experiments in a controlled environment laboratory at the Institut für Biologie–Neurobiologie (Freire Universität Berlin) from February to April 2017. Behavioural assays were conducted in a composed experimental device with two parts, the “arena” (Fig. 1a) and the “dome” (Fig. 1b). The arena was a 30 x 30 x 10 cm empty methacrylate rectangular prism with no roof, containing a grey cardboard as floor and sustained over a wood structure. The dome was a dark brown upside-down plastic flowerpot, illuminated homogeneously with attached LED lamps. The dome covered the arena to create a controlled environment for the experiments. We attached different geometrical figures patterns in the inside walls to facilitate the orientation of the bees during the tests (Jin et *al.* 2014). The dome had a hole in the roof to attach a video camera to record the tests. Citral odour was perfused evenly and restored regularly, as it is known to stimulate bumblebees, and probably other bees, during foraging (Lunau, 1991; Shearer & Boch, 1966).

**Figure 1.**
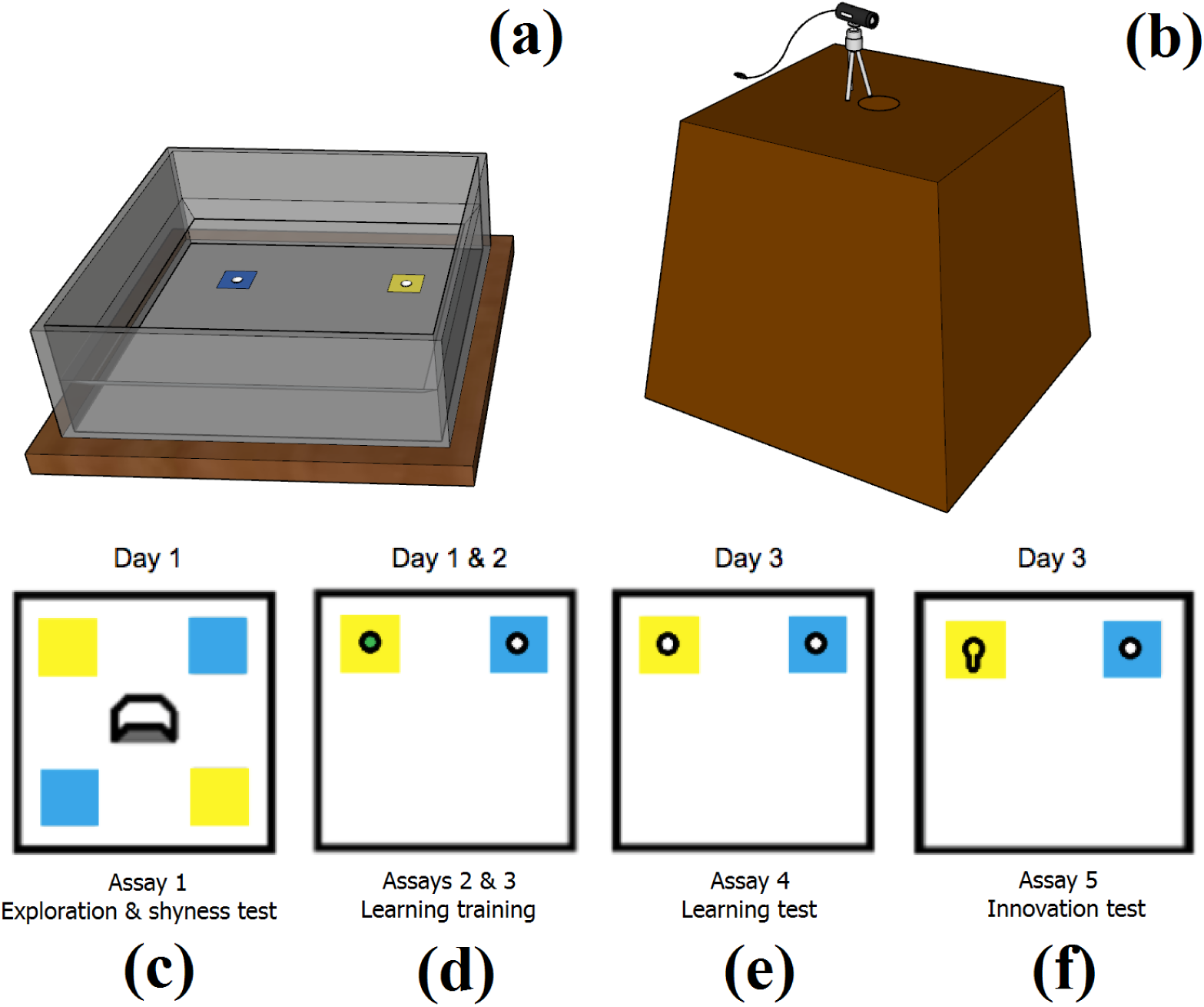
The experimental arena (a) laying in a neutral grey ground, surrounded by plastic walls with plastic cornices attached to avoid that bees can escape. It was covered by the dome (b) with a landscape pattern displayed inside and a webcam placed in the ceiling to record all the experiments. The experiment had four different displays. In assay 1 (c) the bee started inside a refuge. The aim of the assay was to see whether the bee stayed in the refuge (as shyness proxy), and/or explored the colour cues around. In assays 2 and 3 (d), the bee was exposed to two sprues, one rewarded and the other was empty. The colour was randomly selected but maintained along the assays. In assay 4, the learning test (e), the display was the same as in assay 2 and 3, but this time we removed the reward and both sprues were empty. In assay 5, the innovation test (f), the display was the same than in assay 2 and 3 as well, but this time we covered the reward with a lid, forcing the bee to innovate to lift the lid to access the reward.

### Experimental protocol

Along 3 days, each individual passed a sequence of 5 behavioural assays (Fig.1 c, d, e, f) of 15 minutes each designed to measure five different behaviours: exploration, shyness, activity, learning and innovation (see Table 1). Because the mechanisms behind innovation are complex and we do not know what may be driving innovation, we controlled this other related behaviours. We waited four hours between trials if the next trial was done the same day and around 16 if the next trial needed to be done the next day (Fig. 1 c, d, e, f). Activity, measured as the proportion of time in movement, was measured for every trial. Individuals did not show any correlation in their activity levels along the trials (Figure S1) and therefore, we did not estimate a single average activity value for each individual. Activity levels did not decrease along the trials (Linear model Activity ∼ Trial, Estimate ± SE = 0.003 ± 0.008, p = 0.718). Note that not every bee survived to perform all the assays; only 45% of the individuals that started the experiment reached the final assay. Although individuals were not fed during the experimental process other than during the trials, the lack of correlation between the number of feeding events and activity rates during the leaning test (Pearson correlation = −0.09) or the innovation test (Pearson correlation = −0.01) suggests that this high mortality is not attributable to starving.

**Table 1.**
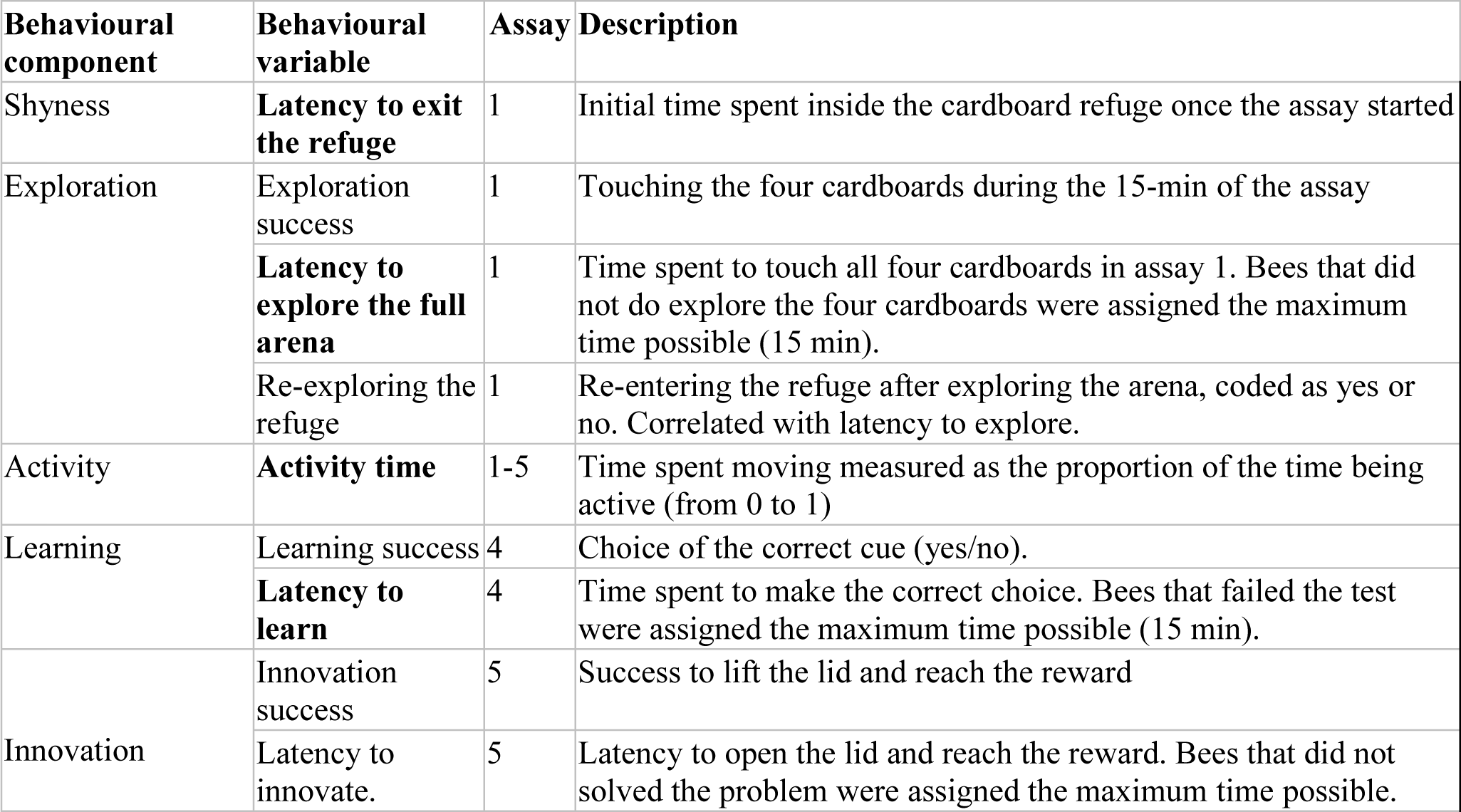
This table contains all variables measured during the tests, with those selected for the innovation analyses as predictors in bold.

| **Behavioural component** | **Behavioural variable** | **Assay** | **Description** |
| --- | --- | --- | --- |
| Shyness | **Latency to exit the refuge** | 1 | Initial time spent inside the cardboard refuge once the assay started |
| Exploration | Exploration success | 1 | Touching the four cardboards during the 15-min of the assay |
|  | **Latency to explore the full arena** | 1 | Time spent to touch all four cardboards in assay 1. Bees that did not do explore the four cardboards were assigned the maximum time possible (15 min). |
|  | Re-exploring the refuge | 1 | Re-entering the refuge after exploring the arena, coded as yes or no. Correlated with latency to explore. |
| Activity | **Activity time** | 1-5 | Time spent moving measured as the proportion of the time being active (from 0 to 1) |
| Learning | Learning success | 4 | Choice of the correct cue (yes/no). |
|  | **Latency to learn** | 4 | Time spent to make the correct choice. Bees that failed the test were assigned the maximum time possible (15 min). |
| Innovation | Innovation success | 5 | Success to lift the lid and reach the reward |
|  | Latency to innovate. | 5 | Latency to open the lid and reach the reward. Bees that did not solved the problem were assigned the maximum time possible. |

The first assay aimed at measuring exploration and shyness. The arena included four coloured cardboard cues (2 blue and 2 yellow, Fig. 1c). The bee was placed in a little cardboard refuge and was kept inside for 5 minutes to allow habituation. Next, the refuge was opened and the individual was allowed to explore the arena. To quantify exploration, we recorded whether the bee explored all the cardboards during the assay and the time it took to do so. Shyness was measured as the initial time spent inside the refuge (Table 1). Re-entering the refuge was originally thought to be a descriptor of shyness, however the analysis of the videos showed that bees did not re-enter the refuge to stay inside and hide, but rather did it as part of their arena exploration

The second and third assays were the learning assays, where we trained bees to associate a colour with a reward (Fig. 1d). The individuals started all tests inside a black opaque box cover that was lifted at the start of the experiment. We displayed 2 cardboards cues with sprues on it, one rewarded with 50% sucrose solution and the other empty. Blue and yellow cardboards are well discriminated by bees (Vorobyev et *al*., 1999; Hempel de Ibarra et *al*., 2014). Hence, the reward for each individual was randomly assigned to one of this two colours for both trials and we let the individuals explore the sprues and eat *ad libitum* during 15 minutes. The position (left or right) of the reward was randomly assigned for each individual in each trial.

In the fourth assay, the learning test, we tested if individuals had learned to associate colours with rewards as trained. The test consisted of both cues displayed as in the second and third assays, but this time with both sprues empty (Fig. 1e). We measured if the individuals approached the formerly rewarded coloured cue and quantified the time spent until checking the right feeder. To ensure that bees had learned to associate colour and reward, we switched the colour of the rewarded sprue for some bees between the two learning assays in 36 randomly selected individuals (control group, hereafter).

In the final assay, we measured the propensity for innovation by using the same coloured cue and reward combination as in assays 2 and 3, but this time the sprue containing the reward was covered with a cardboard lid (Fig. 1f). Bees had thus to innovate -i.e. lift the cardboard-to reach the reward. Innovation propensity was measured in terms of innovation success and latency to succeed (Table 1). Control bees used in the learning assays were not tested for innovation.

### Data analysis

We modelled problem solving performance in the innovation assay as a function of learning, shyness, exploration and activity (see Table 1 for definitions). We modelled the success or failure in solving the task using a Bayesian generalized linear model with a Bernoulli family and a logit link (Package brms; Bürkner, 2017). To model the latency to solve the task, we instead used survival analyses based on cox proportional hazards regressions for continuous predictors (Cox, 2018, Table 2). Survival analysis allow us to add censored data for those individuals that did not passed the test.

**Table 2.**
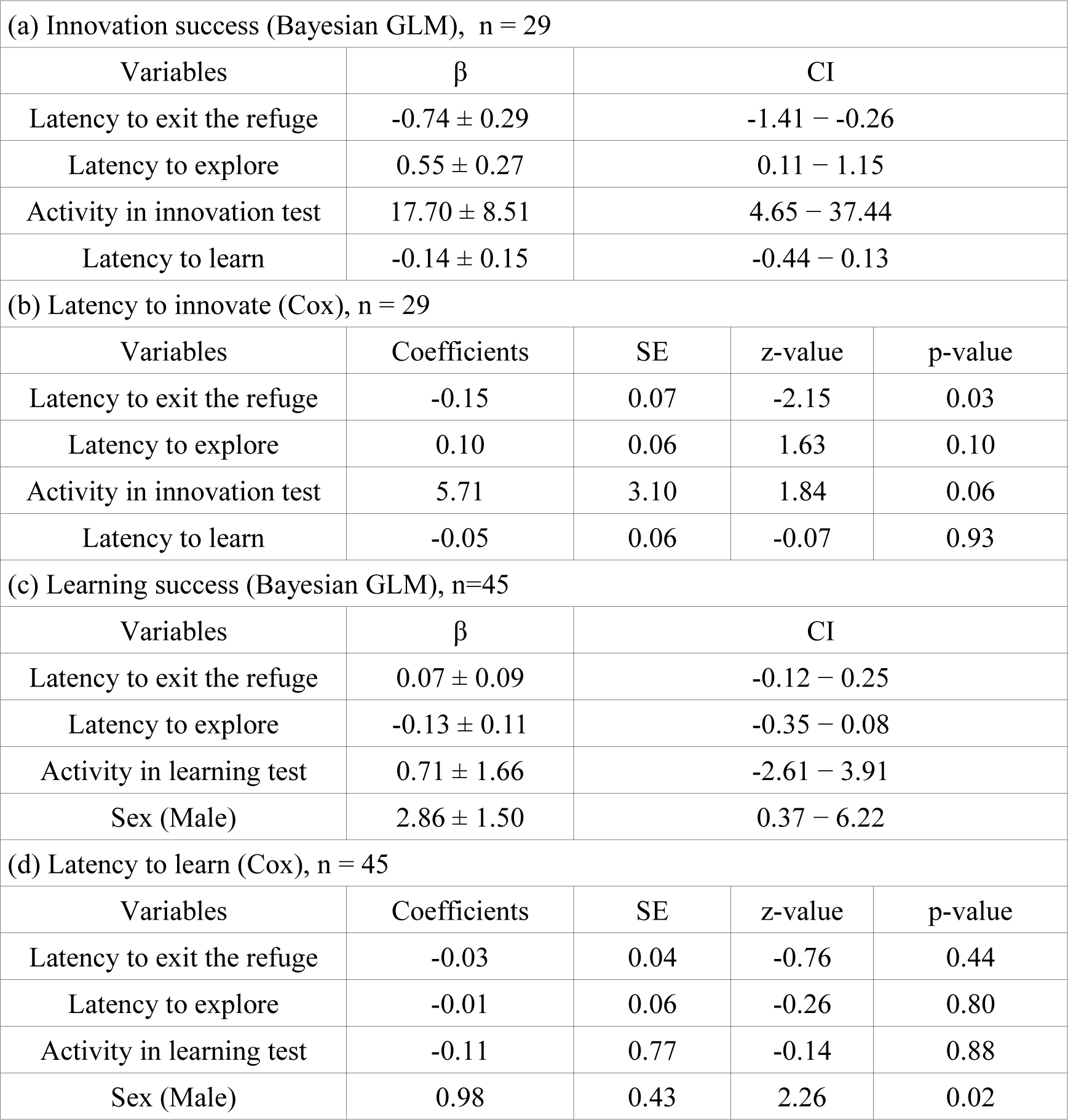
Multivariate model coefficients (beta ± standard deviation) for innovation success and learning as a function of latency learning, shyness, exploration and activity. We ran parallel models for innovation and learning success (Bayesian GLM), and for latency to innovate and learn (Cox). Abbreviations: CI = Confidence interval, Rhat is the potential scale reduction factor on split chains (all our models are at convergence, Rhat = 1).

| (a) Innovation success (Bayesian GLM), n = 29 |  |  |  |  |
| --- | --- | --- | --- | --- |
| Variables | $\beta$ | CI | | |
| Latency to exit the refuge | -0.74 ± 0.29 | -1.41 – -0.26 | | |
| Latency to explore | 0.55 ± 0.27 | 0.11 – 1.15 | | |
| Activity in innovation test | 17.70 ± 8.51 | 4.65 – 37.44 | | |
| Latency to learn | -0.14 ± 0.15 | -0.44 – 0.13 | | |
| (b) Latency to innovate (Cox), n = 29 |  |  |  |  |
| Variables | Coefficients | SE | z-value | p-value |
| Latency to exit the refuge | -0.15 | 0.07 | -2.15 | 0.03 |
| Latency to explore | 0.10 | 0.06 | 1.63 | 0.10 |
| Activity in innovation test | 5.71 | 3.10 | 1.84 | 0.06 |
| Latency to learn | -0.05 | 0.06 | -0.07 | 0.93 |
| (c) Learning success (Bayesian GLM), n=45 |  |  |  |  |
| Variables | $\beta$ | CI | | |
| Latency to exit the refuge | 0.07 ± 0.09 | -0.12 – 0.25 | | |
| Latency to explore | -0.13 ± 0.11 | -0.35 – 0.08 | | |
| Activity in learning test | 0.71 ± 1.66 | -2.61 – 3.91 | | |
| Sex (Male) | 2.86 ± 1.50 | 0.37 – 6.22 | | |
| (d) Latency to learn (Cox), n = 45 |  |  |  |  |
| Variables | Coefficients | SE | z-value | p-value |
| Latency to exit the refuge | -0.03 | 0.04 | -0.76 | 0.44 |
| Latency to explore | -0.01 | 0.06 | -0.26 | 0.80 |
| Activity in learning test | -0.11 | 0.77 | -0.14 | 0.88 |
| Sex (Male) | 0.98 | 0.43 | 2.26 | 0.02 |

In order to avoid model over-parametrization, we used only the quantitative proxies of shyness, exploration and learning (i.e. latencies; Table 1). In addition, as activity levels were variable across trials (Fig. S1), we only included activity levels during the test evaluated. Sex was not added as co-variable because of the limited sample size and skewed proportion of females (6 males, 23 females). Learning success and latency was modelled in a similar way as innovation, that is, as a function of shyness, exploration, activity during the learning test, but this time including sex (9 males, 34 females). For individuals not solving a particular task (e.g. exploration or learning), we assigned to them a maximum latency of 15 minutes.

In summary, for innovation we built multivariate models with latency to exit the refuge (i.e. shyness), latency to explore the full arena (i.e. exploration), latency to perform the learning test (i.e learning) and activity as predictors. For learning we built multivariate models with latency to exit the refuge (i.e. shyness), latency to explore the full arena (i.e. exploration), activity and sex.

## RESULTS

Our experiments showed that *Osmia cornuta* bees were able to innovate. Eleven out of the 29 bees we tested for innovation solved the innovation task, lifting the lid to reach the reward within the 15 minutes of the assay. *Osmia cornuta* bees were also able to learn, with 63% of individuals succeeding in the learning test (n = 48, chi-squared = 3, df = 1, p-value = 0.08) while control bees had a success rate close to that expected by random (n = 36, 52% success, chi-squared = 0.11, df = 1, p = 0.74). Males tended to learn better than females, showing slightly higher success rates (Table 2c) and learning faster (Table 2d). However, latency to innovate showed no relationship with learning (Table 2b, Figure 2b).

**Figure 2.**
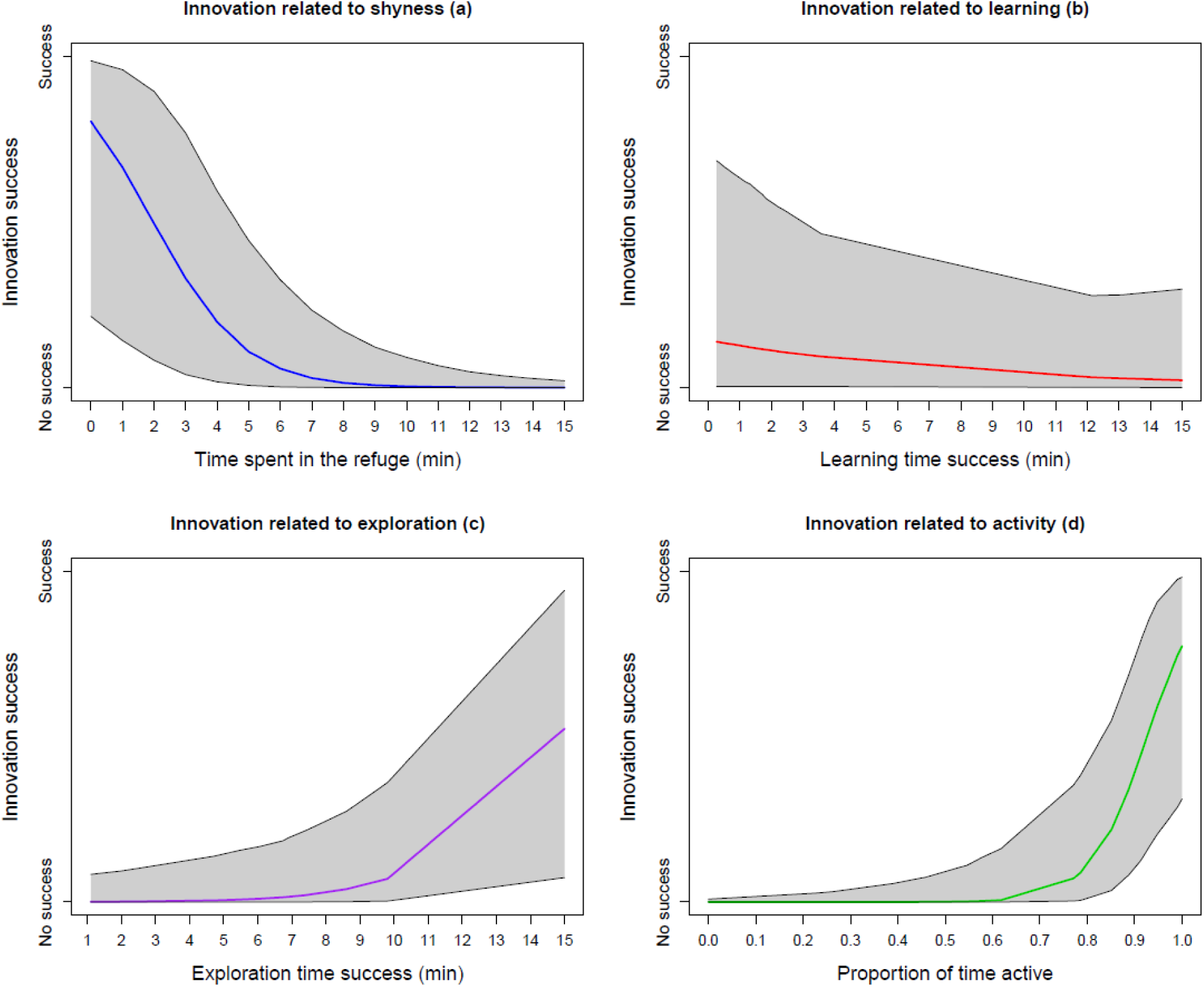
Innovation related to each measured behaviour. These graphs plot the estimates extracted from the multivariate model described in Table 2a measuring the success or failure in the innovation test.

Instead, innovation success and latency were better explained by individual differences in shyness, exploration and activity (Figure 2, Table 2). First, shier individuals were worst innovators. The probability of innovating dropped from 0.80 for bees that spent 2 seconds inside the refuge to 0.01 for bees that did not leave the refuge in the first assay (Table 2a, Fig. 2a). Shier individuals were also slower at resolving the innovation test (Table 2b). In fact, from all bees that did not leave the refuge in the first test (our proxy of shyness) and reached the innovation test, none of them passed the innovation test in subsequent assays.

Second, slower explorers were also better at the innovation test. Bees that spent more time solving the exploration test had more chances to succeed in the innovation test (Table 2a, Figure 2c). These individuals also solved the innovation test faster (Table 2b). Finally, active bees during the innovation test had better chances of solving the innovation test (Table 2a, Figure 2d), indicating that the velocity at solving the test correlated positively with the proportion of time active during the test (Table 2b). Unlike innovation, learning was not affected by shyness, exploration and activity (Table 2b, c; figure 3).

**Figure 3.**
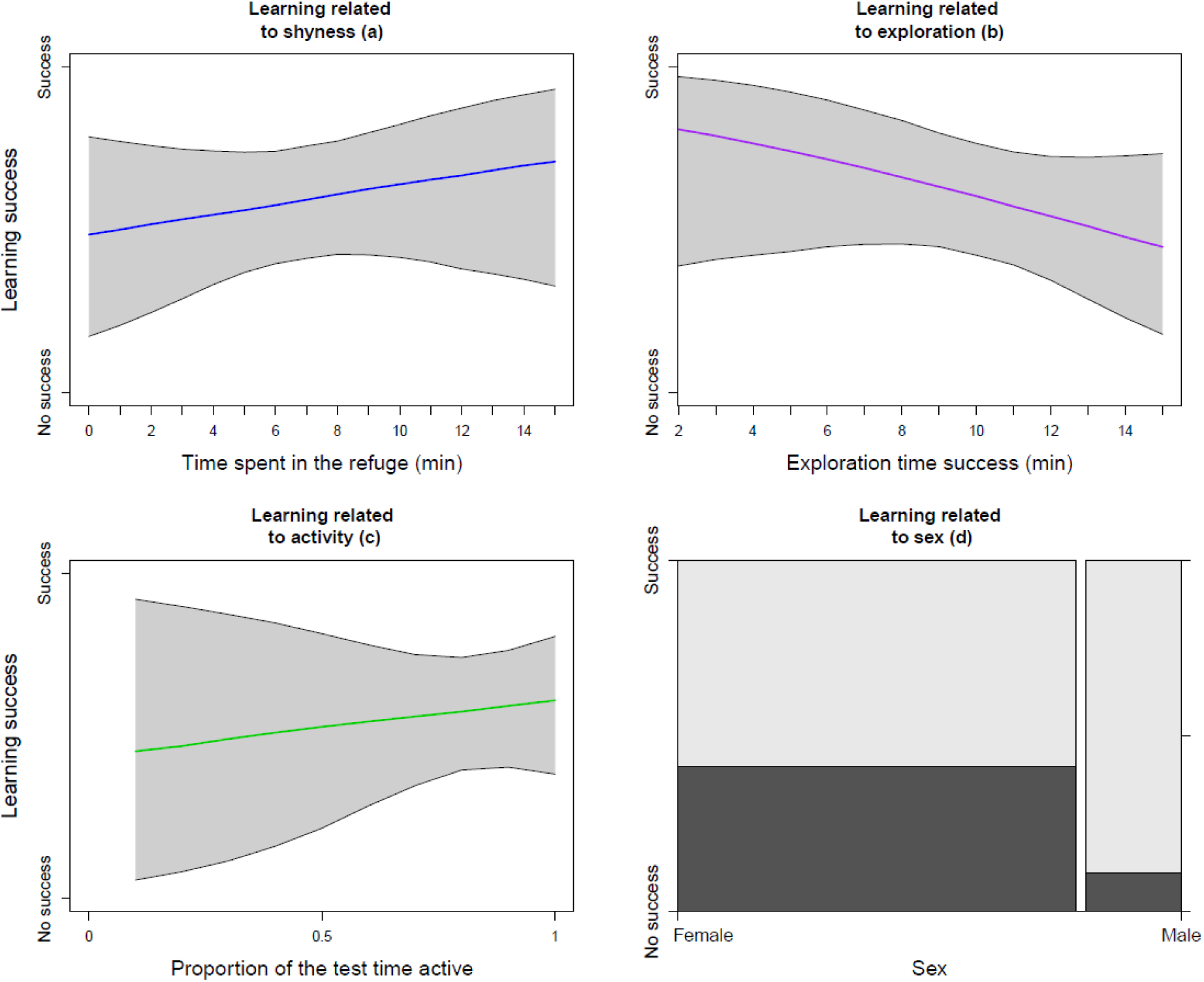
Learning related to each measured behaviour. These graphs are extracted from the multivariate model described in Table 2c measuring the success or failure in the learning test. The width of the bars in (d) is proportional to the number of individuals tested.

## DISCUSSION

Innovation-like behaviours have been previously observed in wild solitary bees. These include the use of new materials for nesting (Allasino et *al*., 2019) and anecdotal examples of bees nesting in new places, such as cardboard, wooden blocks (Bosch & Kemp, 2001) or Styrofoam blocks (MacIvor & Moore, 2013). However, the innovative ability of solitary bees had never been demonstrated before in controlled laboratory experiments. Ours is the first experimental demonstration that *Osmia cornuta* can develop innovative behaviours to solve novel problems.

Although innovation is generally believed to be a dimension of domain-general cognition (Lefebvre et *al.*, 2004), we did not find evidence that individuals the were better at associative learning solved the innovation task faster. The failure to relate innovation and associative learning does not simply reflect that we studied learning over shorter training periods as success in the learning test was comparable to those found in previous similar experiments using more training days (e.g. Jin et *al.*, 2014: Jin et *al.*, 2015).

A more likely explanation is that other factors are more relevant to innovate and can have masked the effect of learning. Indeed, we found consistent differences between fast and slow innovators in their tendency to approach and explore the experimental apparatus. Specifically, individuals that were able to lift the lid to access the food reward tended to be bolder and to explore slower than those that failed to solve the task. As suggested for other taxa, there may be a trade-off between exploration speed and accuracy which can translate into how information is processed. For example, in great tits (*Parus major*), fast explorers return more quickly to previously experienced foraging patches whereas slow explorers prefer to seek new information or update old information close to the feeders (Matthysen et *al*., 2010). Boldness and exploration have been previously identified as important determinants of innovation propensity in vertebrates and highlight that innovation propensity may largely reflect particular motivational states or emotional responses of individuals to novel situations rather than cognitive differences (Sol et *al.*, 2013). In line with this conclusion, successful innovators also exhibited higher activity levels. Activity may reflect motivation to feed, which in other animals has been found to be a major determinant of innovation propensity (e.g. Sol et *al.*, 2013. However, it may also increase the chances to solve the task accidentally by trial and error. Closed environmental spaces can also be stressful and what we defined as “fast exploring” can be a by-product of stereotyped stress behaviours.

The lack of evidence for domain-general cognition does not mean that innovation does not require learning. Learning is not only necessary to fix the new behaviour in the individual repertoire (Ramsey et *al.*, 2007, Lefebvre et *al.*, 2004, Reader et *al.*, 2003, Sol 2003), but it is also important to solve the task itself. Indeed, we found that bees that succeeded in the innovation test went directly towards the lid covering the reward, probably reflecting that they had learnt the rewarding colour during training assays. In our assays, most individuals were able to rapidly associate colours and rewards — after only two training trials— regardless of their differences in shyness, exploration and activity. Thus, the lack of effect of learning ability on innovation might reflect that most individuals were similarly proficient in associative learning.

Learning is widely-held to have important advantages in the wild. In bees, learning is critically important for vital tasks such as foraging, identification of high quality foraging sites, finding the right mixtures of nectar and pollen, and navigating back to the nest for brood provisioning (Roulston & Goodell, 2011; Minckley et *al*., 2013). Surprisingly, we found intriguing sex-related differences in learning. Males showed a tendency to perform better in the associative learning test than females. This is unexpected because females have to deal with more tasks during their lifetime, including foraging and nest provisioning, and may perhaps indicate that the cognitive demands for males to locate females are higher than suspected.

Our results suggest that solitary bees can also readily accommodate their behaviour to novel context through innovative behaviours, with no need of sophisticated cognitive processes. In a context of global change, the ability to rapidly accommodate behaviour to novel contexts seems highly relevant. In novel environments, bees must for instance learn how to forage on new plant species, which sometimes presents complex flowers with whom bees have not co-evolved (Bartomeus et *al.*, 2010). Therefore, we should abandon the notion that insect behaviour is inflexible and stereotypical, and better appreciate that insects can readily accommodate their behaviour to changing conditions through innovation and learning.

## ACKNOWLEDGEMENTS

We wanted to thank Nanxiang Jin for his help on setting the experimental device and the FU Berlin Institute of Biology, for sharing their equipment

## DATA ACCESSIBILITY

The data used for this research will be archived in dryad/figshare upon acceptance. Code used to reproduce the analysis can be consulted at GitHub https://github.com/MiguelAngelCollado/fuocornuta

## AUTHORS’ CONTRIBUTION

MAC, IB and RM designed the experiment; MAC carried out the experimental process under RM supervision; MAC watched the recorded videos from the experiment and extracted the data and wrote the initial draft; MAC and IB did the data analysis with help from DS; All authors contributed to the final version of the article.

## SUPPLEMENTARY MATERIAL

**Figure S1.**
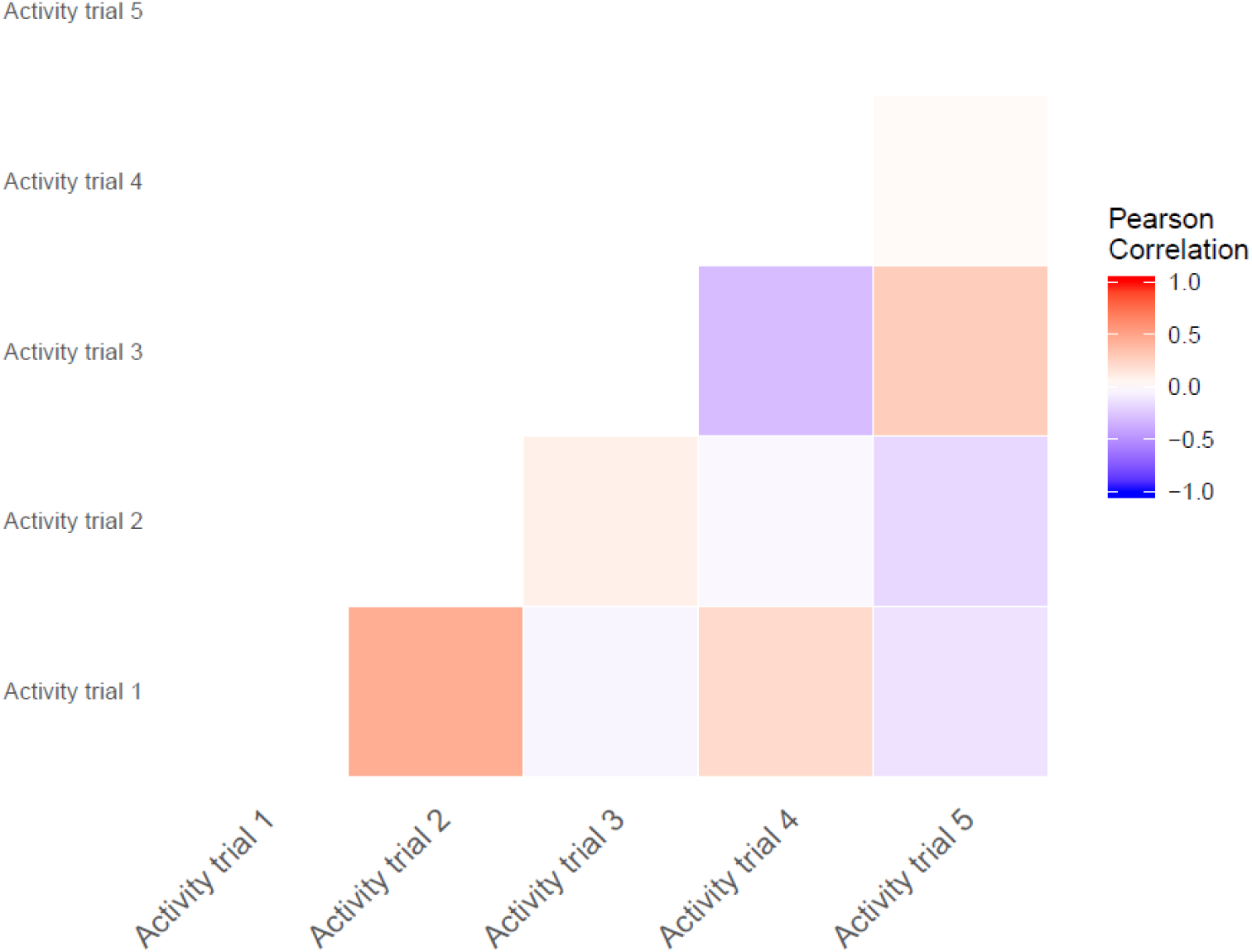
Activity levels across trials, measured as time active, were not correlated (mean Pearson r = 0.23).

