## Supplementary material figure 1 for "Innovation in solitary bees is driven by exploration, shyness and activity levels"

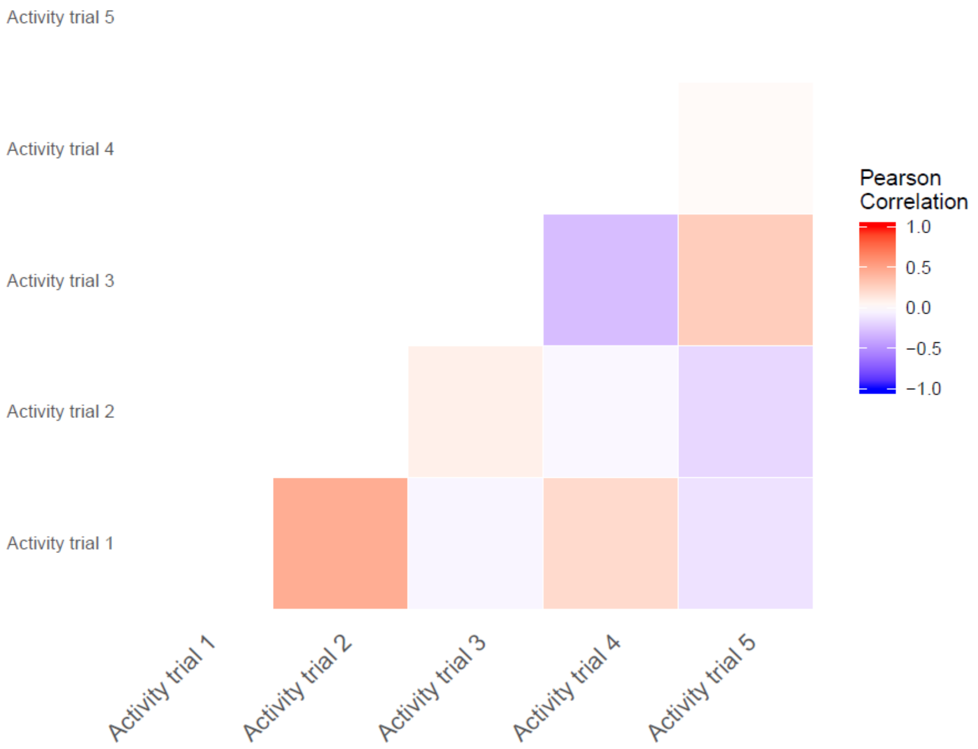


**Figure S1.** Activity levels across trials, measured as time active, were not correlated (mean Pearson r = 0.23).
